# PDX1 directs a core developmentally and evolutionarily conserved gene program in the pancreatic islet

**DOI:** 10.1101/2021.02.28.433241

**Authors:** Xiaodun Yang, Jeffrey C. Raum, Junil Kim, Reynold Yu, Juxiang Yang, Gabriella Rice, Changhong Li, Kyoung-Jae Won, Doris A. Stoffers, Diana E. Stanescu

**Affiliations:** Institute of Diabetes, Obesity and Metabolism, Perelman School of Medicine, University of Pennsylvania, Philadelphia, PA; Department of Medicine, Perelman School of Medicine, University of Pennsylvania, Philadelphia, PA; Biotech Research & Innovation Centre, University of Copenhagen, Denmark; Division of Endocrinology and Diabetes, The Children’s Hospital of Philadelphia, Philadelphia, PA; Department of Cell and Developmental Biology, Institute for Regenerative Medicine, Perelman School of Medicine, University of Pennsylvania, Philadelphia, PA; Department of Pediatrics, Perelman School of Medicine, University of Pennsylvania, Philadelphia, PA

**Author notes:** co-corresponding authors and senior authors (contact and). Co-first authors (Xiaodun Yang, Jeffrey C. Raum).

**Keywords:** Pancreas development, ChIP-seq, single-cell RNA-seq, mouse, human, PDX1 cistrome, PDX1 core gene program

## Abstract

*Pancreatic and duodenal homeobox 1 (PDX1)* is crucial for pancreas organogenesis, yet the dynamic changes in PDX1 targets in mouse or human pancreas development have not been examined. We integrated the PDX1 cistrome with cell lineage-specific gene expression in both mouse and human developing pancreas. We identified a core set of developmentally and evolutionarily conserved PDX1 bound genes that reveal the broad multifaceted role of PDX1 in pancreas development. Despite the well-known, dramatic changes in PDX1 function and expression, we showed that PDX1 binding is largely stable from embryonic pancreas to adult islet. This may point towards a dual role of PDX1, activating or repressing the expression of its targets at different ages, dependent on other functionally-congruent or directly-interacting partners. Our work also suggests that PDX1 functions not only in initiating pancreas differentiation, but also as a potential keepsake of the progenitor program in the adult beta cells.

## Introduction

Diabetes is the most common disorder of the endocrine pancreas and impacts the lives of more than 460 million people worldwide (1). Many environmental and genetic factors have been found to contribute to diabetes pathogenesis. Mutations of key transcription factors cause defects in the endocrine pancreas development and impose susceptibility for diabetes (2). One of the well-studied transcription factors and diabetes genes, pancreatic and duodenal homeobox 1 (*PDX1*), plays essential roles in pancreas organogenesis, endocrine pancreas development, and in the growth and function of insulin-secreting beta cells in both mouse and human (3, 4).

PDX1 is the hallmark transcription factor of pancreas development. Pancreas organogenesis starts upon the induction of *Pdx1*/*PDX1* in the foregut endoderm at embryonic day 8.5 (E8.5) in mouse (5) and at approximately 29 days post conception (dpc) in human (6). In mouse, PDX1+ pancreatic progenitors proliferate to expand from E9.5 to E12.5 (7). PDX1 is then gradually enriched in mouse pancreatic beta cells at later stages (reviewed by (3)), but low level PDX1 expression is detected in acinar and ductal cells in the adult mouse pancreas (8). In human pancreas development, PDX1 is expressed in a similar pattern (reviewed by (4)). Homozygous loss-of-function of *Pdx1*/*PDX1* causes pancreas agenesis in both mouse (9, 10) and human (11). Heterozygous mutations in *PDX1* are linked to human type 2 diabetes (12, 13). Over the past 10 years, many studies have contributed to the effort to understand the complex role of PDX1 in both pancreas development and adult beta-cell function. Mouse models with *Pdx1* hypomorphic alleles (14) or heterozygous loss-of-function of *Pdx1* (15) have provided valuable information about the gene networks regulated by PDX1. It interacts with ONECUT1 to directly activate endocrine progenitor gene *Neurog3* expression and coordinates a transcription factor network, including *Foxa2*, *Sox9*, *Onecut1*, *Hnf1b* (14, 15). PDX1 genome-wide target genes have been identified in human embryonic stem cell-derived pancreatic progenitors (16–18) and in adult mouse and human islets (19). It regulates important transcription factors in pancreas development *in vitro* (18) and genes functioning in endocrine system and metabolic disorders in adult islet *in vivo* (19). However, a significant gap of knowledge remains regarding the temporal differences in PDX1 targets in pancreas development and adult islet *in vivo*. Furthermore, although mouse models have been the mainstay of fundamental research in endocrine pancreas development, there is no detailed understanding of the differences between the developmental PDX1 targets in mouse and human.

In order to answer these important questions in the field, we performed PDX1 chromatin immunoprecipitation followed by high throughput sequencing (ChIP-Seq) in human fetal pancreata at 14 weeks gestation and mouse embryonic pancreata at E13.5 and E15.5. We characterized in parallel the transcriptomic signatures of human and mouse pancreas cells and integrated them with the developmental PDX1 cistrome to identify the temporal pattern of PDX1 targets expressed in each lineage. We show that PDX1 bound important acinar and ductal genes, from early development to adult islet in both human and mouse pancreas. The integration of our human and mouse cistromic and transcriptomic data identified a conserved ductal and endocrine program directed by PDX1. These results reveal a core developmentally and evolutionarily conserved PDX1 gene program that establishes pancreatic lineage and maintains pancreatic function and suggest PDX1 as a keepsake of an embryonic pancreatic progenitor program in adult beta cells.

## Materials and Methods

### Animals

Mouse experiments were performed at University of Pennsylvania, with IACUC approval. Timed pregnant CD1 mice were purchased from Charles River Laboratories (Wilmington, MA). Dams were housed in AAALC approved rodent colony. Embryonic pancreata were dissected at E13.5 and E15.5.

### Human fetal pancreas samples

Fetal human pancreas tissues were obtained in the period of time between 2011-2017 from StemExpress (Los Angeles, CA) and from Advance Bioscience Resources (Alameda, CA) after elective termination of pregnancy. An informed consent was obtained at the time of the elective termination of pregnancy and is held by supplying company. A sample of the Advance Bioscience Resource consent was obtained and complies with NIH requirements (specifically donation of human fetal tissues was obtained by someone other than the person who obtained the informed consent for abortion, occurred after the informed consent for abortion, and will not affect the method of abortion; no enticements, benefits, or financial incentives were used at any level of the process to incentivize abortion or the donation of human fetal tissues; and to be signed by both the woman and the person who obtains the informed consent). A total of 7 samples were used for these experiments (3 pancreas samples at 14 weeks gestation, and 1 sample each at 15, 18, 19 and 20 weeks gestation). Gestational age was determined by ultrasound, usually using a crown-rump length. No other information on maternal history was available. Experiments performed on these samples were exempt from IRB evaluation by the University of Pennsylvania and The Children’s Hospital of Philadelphia. Samples were received deidentified, in RPMI/5%FBS/antibiotics media on ice. All samples were processed for ChIP-Seq or single cell RNA-seq the day they were received as detailed below.

### ChIP-seq

We performed ChIP-seq for PDX1 in mouse embryonic pancreata (at E13.5 and E15.5) and in human fetal pancreata (at 14 weeks gestational age). We also performed ChIP-seq for H3K27ac and H3K27me3 marks in mouse pancreata at E13.5 and E15.5. For the Pdx1 chromatin immunoprecipitation, we used the goat anti-PDX1 antibody (gift from Chris Wright). For the histone marks – anti-H3K27ac and anti-H3K27me3 antibodies were used (Abcam, Cambridge, MA). 50-60 individual mouse pancreata were pooled for one chromatin immunoprecipitation replicate. All analysis was performed in 3 replicates. Chromatin was prepared for individual replicates from 3 human fetal pancreata. ChIP-seq was performed as previously described (19). All samples were sequenced in the Functional Genomic Core at University of Pennsylvania. The ChIP-seq reads were mapped to the mouse genome (mm10) using Bowtie.

The PDX1 binding in hPSC-derived pancreas progenitor cells was obtained from previously published data by the Lickert laboratory (18). The PDX1 binding in adult human and mouse islets were previously published by our group (19). After peak calling by HOMER, we continued the analysis with approximately top 25% of peaks. For the mouse data, at e13.5 there were 4107 peaks (lowest HOMER score 5.28), at e15.5 were 5343 peaks (lowest HOMER score 5.03) and in mouse islet were 4559 peaks (lowest HOMER score 5.9) (Supplemental Tables S9-10). For the human data, in fetal pancreas there were 8183 peaks, representing top 25% of the peaks (lowest HOMER score 11.1 – corresponding to 6294 genes), hPSC-PP cells there were 21292 peaks (lowest HOMER score – 18.9), corresponding to 11451 genes, and for human adult islet there were 17593 peaks (lowest HOMER score 18.2) – corresponding to 8307 genes. The majority of PDX1 peaks were within 2000 bp of the transcription start sites (TSS) (Supplemental Figure 1B). Gene ontology analysis was performed with Ingenuity Pathway analysis software (Qiagen, Germantown, MD). BioVenn and DeepVenn were used to generate the Venn diagrams. (20)

### Single-cell RNA-seq (scRNA-seq) and data analysis

Single islet cells were dissociated with TrypLe, and loaded on the 10x Genomics (Pleasanton, CA) platform in the Center for Applied Genomics at The Children’s Hospital of Philadelphia. Single-cell transcriptome libraries were sequenced on the HiSeq platform (Illumina, San Diego, CA). The cellranger (v.2.1,10x genomics) pipeline was used for barcode filtering, alignment (to GRCh38), and UMI counting. Secondary bioinformatic analysis was performed using the Seurat (v.2.3.4) packages in R. Briefly, sequenced 10x libraries were individually evaluated on multiple criteria to determine cells of interest including low expression of Hemoglobin (average log2 UMI of HBA1, HBA2, HBB < 8) and number of UMI (0 – 65536). Two scRNAseq samples of fetal and adult were integrated by anchor-based method implemented in Seurat. The top 30 principal components were used to perform clustering and visualization using a UMAP (Uniform Manifold Approximation and Projection) plot. Gene marker expression was used to identify specific populations. To identify differentially expressed genes between cell populations in fetal and adult cells, a non-parametric Wilcoxon rank sum test was performed, and p-values were adjusted using a Benjamini-Hochberg correction based on total number of genes in the dataset.

Human adult islet cell transcriptomic data were previously published (GSE114297, (21)). Mouse transcriptomic data of the developing pancreas were from (22, 23).

### Immunofluorescence

A small portion of the human fetal pancreas samples at 14 weeks gestation were fixed with 4% PFA overnight, embedded in OCT and stored at −80C. Cryosections were then used for immunofluorescence for PDX1 with goat anti-Pdx1(Santa Cruz Biotechnology, A-17, 1:500), rabbit anti-Onecut1 (Santa Cruz Biotechnology, H-100, 1:1000), and guinea-pig anti-Insulin (Abcam, 1:500).

### Data Availability

All mouse embryonic and human fetal PDX1 ChIP-seq data were made publicly available and uploaded to ArrayExpress (E-MTAB-3354). The human fetal pancreas single-cell RNA-seq data are available upon request from DES.

## Results

### Temporal analysis of PDX1 bound genes in human pancreas development and adult islets

To examine PDX1 genome-wide targets in human developing pancreas, we performed PDX1 ChIP-seq using human fetal pancreata at 14 weeks gestational age. HOMER *de novo* motif analysis showed that the most enriched binding motif matched consensus PDX1 binding motif, indicating that our PDX1 ChIP-seq data were efficient and specific (Supplemental Figure 1A). All fetal pancreatic cells expressed PDX1 at 14 weeks gestation, including insulin positive beta cells and non-hormone positive cells labelled by ONECUT1 (Supplemental Figure 2). We compared this newly generated PDX1 cistromic data (Supplemental Table S1) with published PDX1 binding in human stem cell-derived pancreatic progenitors (hPSC-PP) (Supplemental Table S2) (18) and adult islets (Supplemental Table S3) (19). A total of 3131 genes were bound by PDX1 in all 3 data sets (Figure 1A). These represented approximately 50% of the human fetal pancreas bound genes. Only 1092 and 1826 genes were bound by PDX1 only at fetal or adult stages, respectively (Figure 1A). These data indicated that PDX1 binding is largely preserved from fetal to adult stages.

**Figure 1:**
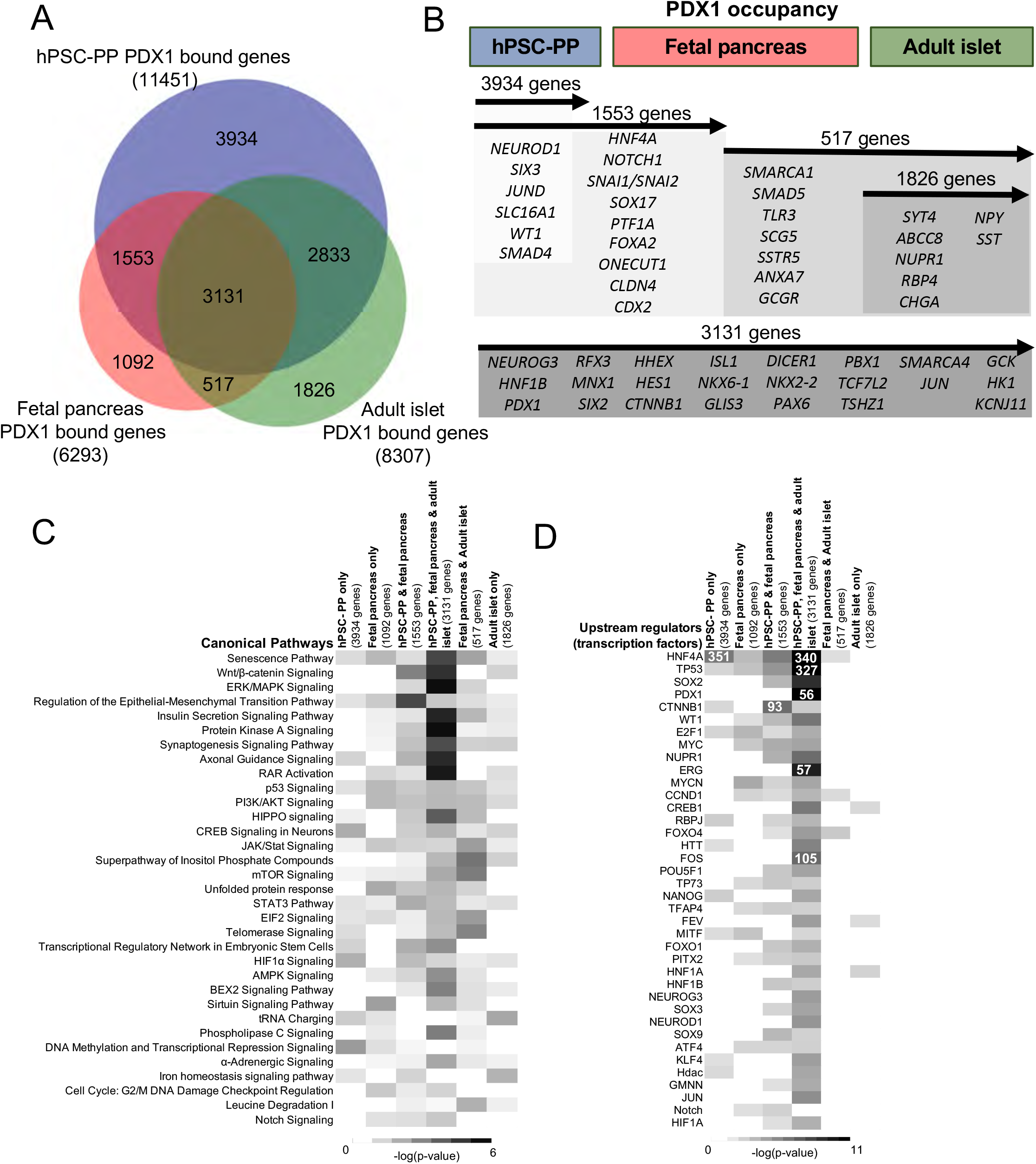
Temporal analysis of PDX1 binding in human pancreas development and adult islets. A. Venn diagram of PDX1 bound genes in hPSC-PP cells, fetal pancreas and adult islet. B. Temporal arrangement of representative genes bound by PDX1 in hPSC-PP, fetal pancreas and adult islet. C. Gene ontology analysis of canonical pathways enriched in subsets of PDX1 bound genes at developmental and adult stages. D. Upstream regulator analysis of transcription factors predicted to directly regulate PDX1 bound genes at developmental and adult stages.

We arranged these data sets based on their developmental stage, in order to analyze the temporal changes in PDX1 binding. We considered hPSC-PP target binding as an earlier developmental stage than the fetal pancreas binding. This allowed a timeline comparison of PDX1 bound targets from an early developmental stage (hPSC-PP), to the fetal pancreas and in adult islet (Figure 1B). We identified genes that were bound only in pancreas progenitors (*NEUROD1*, *JUND*, *SIX3*), genes that were bound only during development (*HNF4A*, *ONECUT1*, *CLDN4*, *PTF1A*) and genes that were bound only in adult islet (*SYT4*, *NPY*, *CHGA*) (Figure 1B). Significantly, PDX1 continuously bound genes with important roles in the specification and differentiation of pancreatic endocrine progenitors (*NEUROG3*, *NKX6-1*, *NKX2-2*, *RFX3*, *HNF1B*) and/or in the function of adult beta cells (*JUN*, *GCK*, *KCNJ11*). This supported a temporally versatile role of PDX1 in coordinating individual targets.

We compared gene ontology categories for PDX1 bound genes for each subset in order to further clarified the temporal roles of PDX1. Even if the bound genes were distinct, several key signaling pathways appeared to be consistently coordinated by PDX1 across all subsets, including the *senescence pathway, WNT/beta-catenin signaling*, *epithelial-mesenchymal transition*, *insulin secretion signaling pathway* and *protein kinase A pathway*. Several pathways appeared enriched only in the developmental subsets (hPSC-PP or fetal pancreas), such as the *axonal guidance signaling, mTOR signaling, Notch signaling.* Pathways enriched only in the adult islets were *tRNA charging*, *iron homeostasis* (Figure 1C, Supplemental Table S4). We then compared predicted upstream regulators for each of the subsets of PDX1 bound genes in Ingenuity IPA, in order to identify the potential PDX1-interacting protein partners. A large number of PDX1 bound genes in human pancreas development are also known direct targets of transcriptions factors with known roles in pancreas development (HNF4A, SOX2, beta catenin, HNF1A, NEUROD1) (Figure 1D, Supplemental Table S5). Surprisingly, we found p53 (*TP53*) as the second predicted upstream regulator of developmental PDX1 bound genes. Although p53 has clear roles in pancreatic cancer pathogenesis (24, 25) and may affect pancreatic progenitor cell survival (26), its role in pancreas development has not yet been well investigated. These findings expand the list of transcription factors and cofactors that are potentially either functionally-congruent or directly-interacting PDX1 partners.

### Comparative single-cell transcriptome analysis of human fetal pancreas and adult islet cells reveals cell-lineage-specific and temporally regulated genes

The developing pancreas is composed of multiple cell types at different stages of differentiation. In order to understand the role of PDX1 in each cell lineage, we performed single-cell RNA-seq analysis of human fetal pancreas at 15-20 weeks gestational age. We analyzed these newly generated fetal cell transcriptomes together with adult pancreas cell transcriptomes (21). A total number of 7,421 fetal cells and 20,764 adult pancreatic cells were examined. We assessed cell-type-specific gene expression signatures of acinar, ductal, and endocrine cells (Figure 2A-B). Using cell lineage specific markers, we identified pancreatic cell clusters for alpha cells (*GCG*), beta cells (*INS*), delta cells (*SST*), PP cells (*PPY*), acinar cells (*PRSS1*), and ductal cells (*CFTR*) (Figure 2A, Supplemental Figure 3). Fetal and adult pancreatic cells largely overlapped in each cluster (Figure 2A-B). A significant number of genes were differentially expressed among fetal and adult cells of the same lineage (Supplemental Figure 4A-F). These differences in gene profiles between fetal and adult pancreatic cells constitute a resource for further investigation of maturational changes in pancreas development. Based on the differentially expressed genes within each age (fetal/adult), we constructed transcriptomic signatures for each cell lineage (Figure 2C-D, Supplemental Table S6, S7).

**Figure 2:**
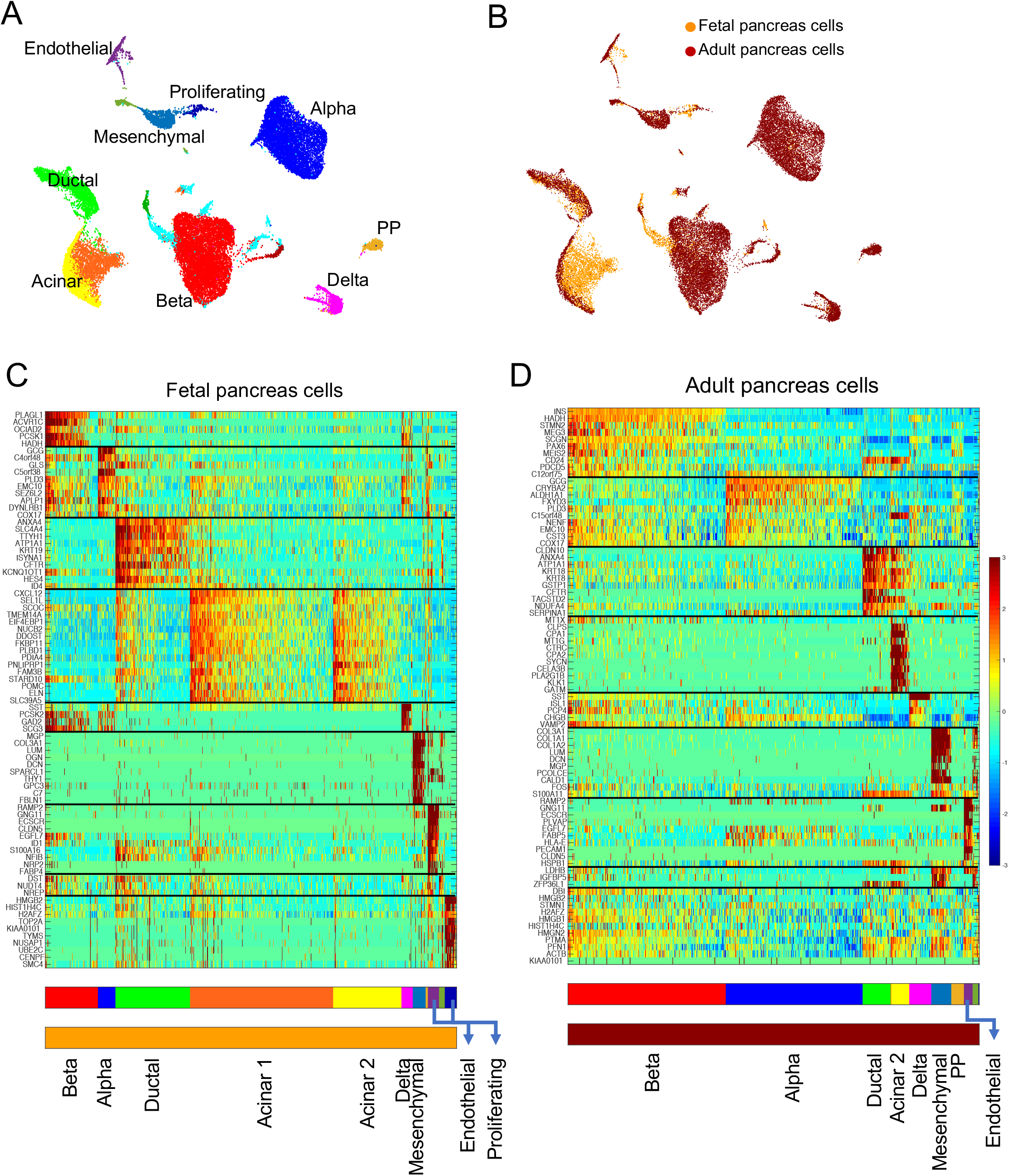
Comparative single-cell transcriptome analysis of human fetal pancreas and adult islet cells reveals cell-lineage-specific and temporally regulated genes. A-B. UMAP plots showed cell clusters (A) identified in single-cell RNA-seq from fetal and adult cells (B). C. Heatmap showed differentially expressed genes in each cluster among human fetal pancreas cells. D. Heatmap showed differentially expressed genes in each cluster among in human adult pancreas cells. For C-D, the DEGs in each cluster were picked up by FDR <0.01 and log ratio > 0.5. The DEGs commonly found in multiple clusters were excluded. If there is more than 10 DEGs in each cluster, only top 10 DEGs by log ratio were included in the heatmap.

### Integrating PDX1 ChIP-seq with single-cell RNA-seq identifies functional roles of PDX1 in regulating its targets in human fetal pancreas and adult islets

We next sought to infer cell-lineage-specific PDX1 targets by integrating single-cell transcriptomics with PDX1 cistrome data in human pancreas. We identified functional PDX1 targets that are upregulated in fetal or adult beta cells, ductal cells, and acinar cells, respectively (Figure 3A, Supplemental Table S8). PDX1 targets enriched in beta cells increased from fetal to adult stages, while those enriched in ductal and acinar cells decreased, indicating a shift of PDX1 binding to maintain beta cell identity and function. The cell-lineage specific PDX1 targets include *ASCL2*, *MEIS2*, *ISL1*, *MAFB* in fetal beta cells, *FXYD2*, *CPA1*, *KRT8*, *TPM4* in fetal ductal cells, *CPA1*, *BTG1*, *KRT8*, *KRT18* in fetal acinar cells (Figure 3B). The comparison between fetal and adult stages revealed stage-specific and conserved signature genes in fetal beta cells, ductal cells, and acinar cells (Figure 3C). For example, *ASCL2* is a target gene of Wnt signaling (27) and expressed in human mature beta cells (28), which was enriched in fetal beta cells and bound by PDX1 only in fetal beta cells. *MEIS2* and *MAFB* were directly regulated by PDX1 in both fetal and adult beta cells. *SNAP25, ABCC8, SCG2* were enriched in the adult beta cells and also bound by PDX1 only in adult islets. PDX1 surprisingly bound important early acinar and ductal specific genes, not only during early development but also in the adult human islet. For example, fetal and adult specific ductal genes, such as *HHEX, ANXA2, KRT8 and KRT18* were bound by PDX1 from fetal pancreas to adult islet (Figure 3C). Similarly, acinar specific genes, such as *CPA1, RBPJ, SPINK1, CELA2A*, were expressed in fetal and/or adult cells and were bound by PDX1 in fetal pancreas and in adult islet. These findings suggest that PDX1 establishes an early developmental binding at acinar and ductal-specific genes, and it surprisingly remains at these targets in the adult islet, despite changes in cellular functions.

**Figure 3:**
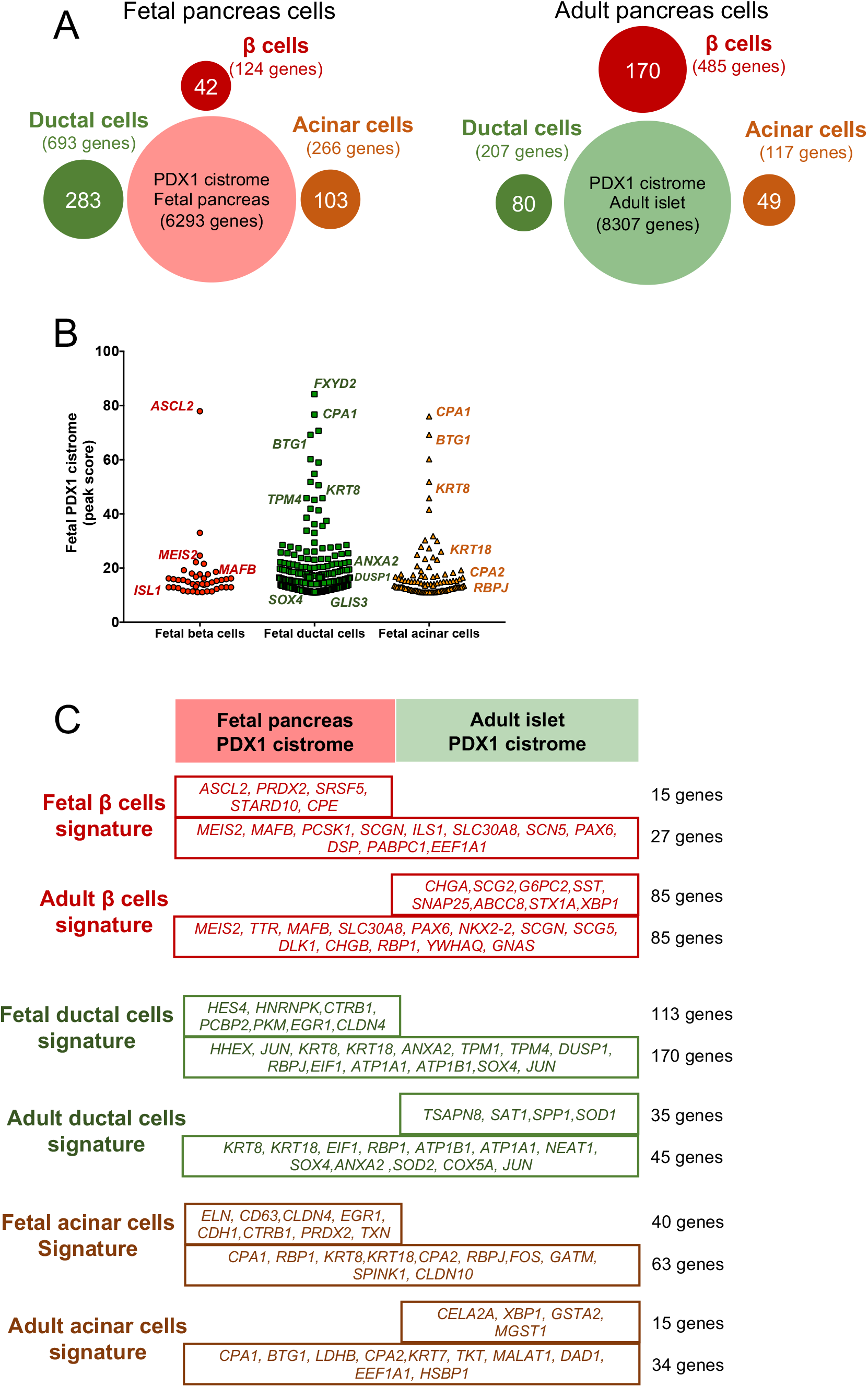
Integrating PDX1 ChIP-seq with single-cell RNA-seq identifies functional roles of PDX1 in regulating targets in human fetal pancreas and adult islets. A. PDX1 bound genes differentially upregulated in human fetal and adult beta, ductal, and acinar cells. B. Peak scores of PDX1 bound genes, expressed in fetal beta, ductal, and acinar cells. C. Stage-specific PDX1 bound genes differentially upregulated in beta, ductal, and acinar cells.

### Temporal analysis in PDX1 binding and cell-lineage-specific target expression from pancreas development to adult islet in mouse

To examine species-conserved and specific PDX1 binding in mouse pancreas, we performed PDX1 ChIP-seq using mouse embryonic pancreata at E13.5 and E15.5 (Supplemental Table S9, S10). We compared this newly generated data with published PDX1 cistrome in mouse adult islets (19) (Supplemental Table S11). We focused on E13.5 because there are a large number of pancreatic progenitors in the mouse developing pancreas after the expansion during the primary transition (E9.5-12.5) (29), which start to lose their multipotency (30). Furthermore, PDX1 is widely expressed in mouse developing pancreas but at high levels in embryonic beta cells at E15.5 (3). The most enriched binding motifs of PDX1 identified at E13.5 (Supplemental Figure 5A) and E15.5 (Supplemental Figure 5B) both matched the PDX1 consensus motif using HOMER, indicating our ChIP-seq experiments were efficient and specific (Supplemental Figure 6).

Similar to the human cistromic analysis, the majority of PDX1 target genes (2040) were continuously bound by PDX1 in mouse pancreas from embryonic to adult stages, while 212, 425, and 389 genes were bound by PDX1 only at E13.5, E15.5, and adult stage, respectively (Figure 4A). Among the developmental specific PDX1 bound genes were *Dlk1, Hnf1b, Pax4, Mki67*. Adult islet specific targets were *Rfx6, Slc30a8, Six3, Scg2* (Figure 4A). Most of the known genes important for pancreas developmental and adult beta cell function were continuously bound from E13.5-E15.5 into the adult islet: *Neurog3, Onecut1, Pax6, Foxs2, Mnx1, Nkx6-1, Rfx3*. Pathway analysis revealed that despite temporal distinct targets, PDX1 regulated key signaling pathways throughout development and in adult islet including the *WNT/beta-catenin signaling*, *epithelial-mesenchymal transition*, and *Protein Kinase A pathways*, which are important for cellular proliferation and/or differentiation (Figure 4B, Supplemental Table S12). Upstream regulator analysis suggested that a large number of these PDX1 targets are co-regulated by other transcriptional factors and cofactors, including p53, SOX2, beta-catenin, and LDB1, similar to the results of the analysis from human pancreas (Supplemental Figure 5C, Supplemental Table S13).

**Figure 4:**
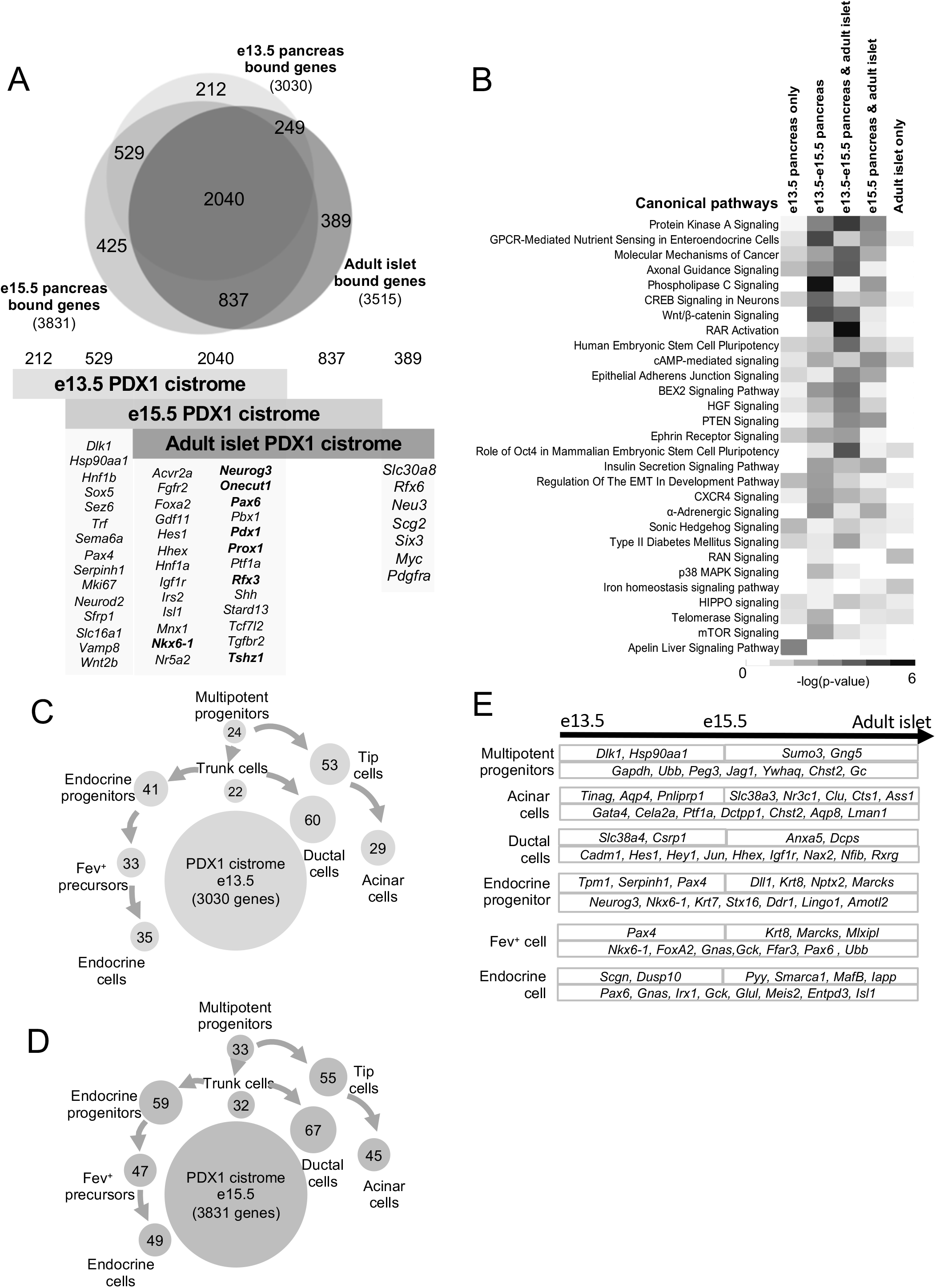
Temporal analysis of PDX1 binding and cell-lineage-specific target expression from pancreas development to adult islet in mouse. A. Venn diagram of PDX1 bound genes in E13.5, E15.5 and adult islet. Representative genes expressed among the 3 data sets (below) B. Gene ontology analysis of canonical pathways enriched in subsets of PDX1 bound genes in the developing mouse pancreas and adult islet. C-D. PDX1 bound genes expressed in different cell types in mouse developing pancreas at E13.5 (C) and E15.5 (D). E. Representative PDX1 bound genes upregulated in different cell types.

In order to characterize the functional roles of PDX1 in different cell types during mouse pancreas development, we integrated the identified PDX1 targets with published single-cell RNA-seq data of mouse developing pancreas (22, 23). We identified PDX1 bound genes that were highly expressed in multipotent progenitors (including *Dlk1*), endocrine progenitors (including *Neurog3*, *Neurod2*), embryonic endocrine cells (including *Pax6*, *Mafb*), and other cell types (Figure 4C-E, Supplemental Figure 5E, Supplemental Table S14, S15). There were more PDX1 targets enriched in the endocrine lineages at E15.5 compared to those at E13.5 (Supplemental Figure 5D), indicating the important roles of PDX1 in endocrine pancreas development at later stages. To examine the functional status of genome-wide promoters and enhancers during mouse pancreas development, we generated H3K27ac (an active histone mark) and H3K27me3 (a repressive histone mark) ChIP-seq data using mouse embryonic pancreata at E13.5 and E15.5 (Supplemental Figure 5F, Supplemental Table S16). We identified a trend of increasing PDX1 binding at active promoters and enhancers from E13.5 to E15.5, indicating the gradual activation of PDX1 targets during pancreas development.

### PDX1 directs a developmentally and evolutionarily conserved ductal and endocrine program

To examine the species-conserved and specific PDX1 targets in mouse and human pancreas, we compared PDX1 bound genes at developmental and adult stages. We found that 762 genes (Figure 5A) and 1509 genes (Figure 5B) were bound by PDX1 in the developing pancreas and adult islets, respectively, in mouse and human. Of these, 518 genes were conserved both developmentally and evolutionarily (Figure 5C, Supplemental Table S17).

**Figure 5:**
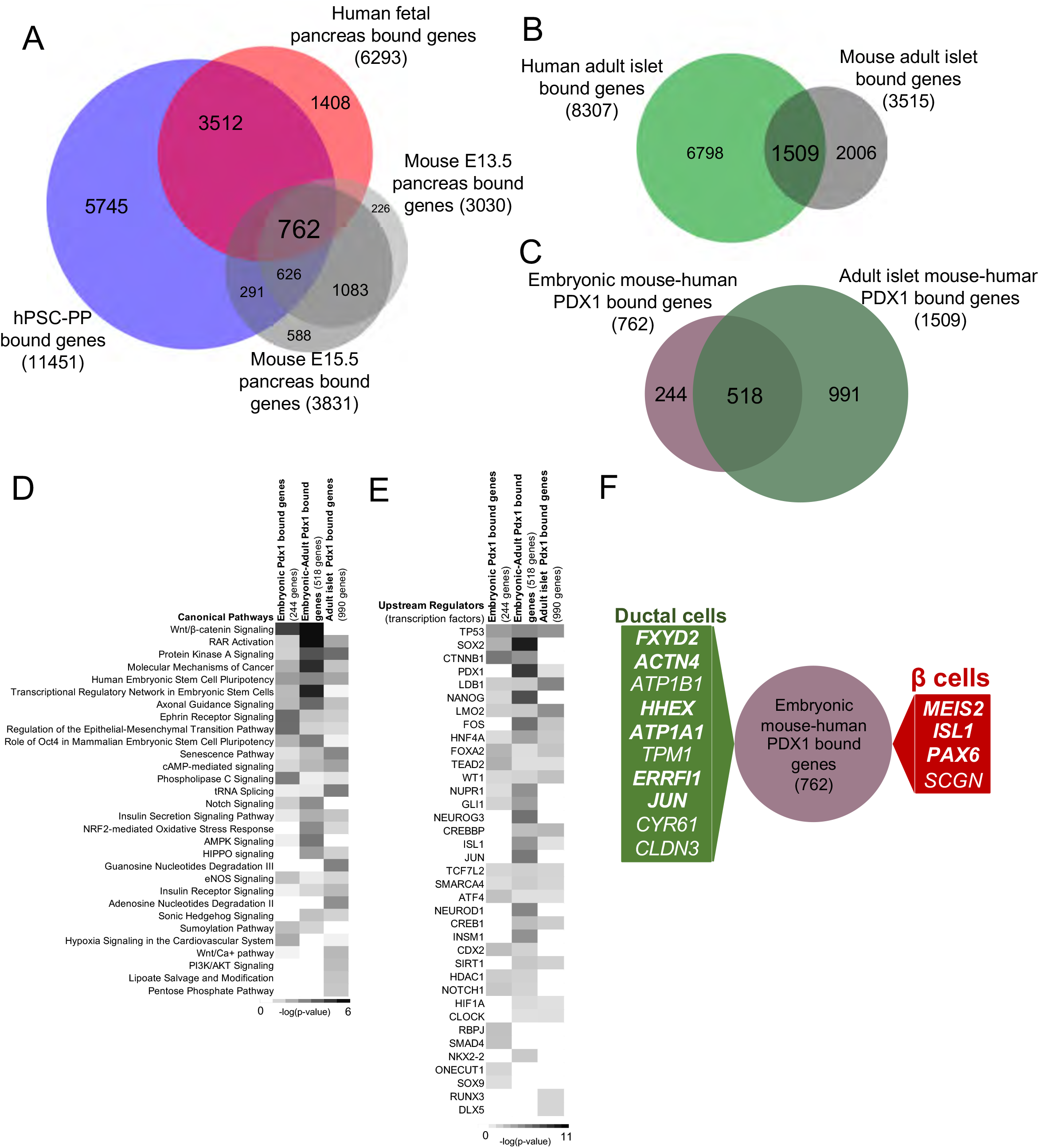
PDX1 directs a developmentally conserved ductal and endocrine program. A. Venn diagram of PDX1 bound genes in human and mouse pancreas development. B. Venn diagram of PDX1 bound genes in human and mouse adult islet. C. Venn diagram of PDX1 bound genes in pancreas development and in adult islet in mouse and human. D. Gene ontology analysis of canonical pathways enriched in subsets of PDX1 bound genes in pancreas development and adult islet in mouse and human. E. Upstream regulator analysis of transcription factors predicted to directly regulate PDX1 bound genes in pancreas development and adult islet in mouse and human. F. PDX1 bound genes expressed in mouse and human fetal ductal or beta cells. The genes in bold are also bound by PDX1 in the mouse and human adult islet.

Pathway analysis of the conserved PDX1 targets showed temporal shifts in key signaling pathways. Several pathways were primarily represented at the developmental stages (*Wnt/beta catenin signaling, Notch signaling, AMPK signaling, role of OCT4 in embryonic stem cell pluripotency*), several were represented only in the adult islet (*Pi3K/AKT signaling, pentose phosphate pathway*), while most appeared in all 3 subsets (*protein kinase A signaling, axonal guidance signaling*) (Figure 5D, Supplemental Table S18). Similar to the mouse and human analyses, several transcription factors were predicted to directly regulate PDX1 bound targets both in embryonic pancreas and in the adult islet. Among these, p53, SOX2, beta catenin, and LDB1 appeared as the top predicted upstream regulators (Figure 5E, Supplemental Table S19). These findings show that PDX1 could physically or functionally interact with many transcription factors and cofactors, and this pattern of interaction is remarkably well conserved between mouse and human.

We next integrated the embryonic mouse-human PDX1 bound genes with the single cell transcriptomic signatures of ductal, acinar and beta cells. None of the genes enriched in human fetal acinar cells or in mouse embryonic acinar cells were part of the embryonic mouse-human PDX1 cistrome. Ten genes were expressed in embryonic ductal cells and were bound by PDX1 in pancreas development in both mouse and human (Figure 5F). Six of these genes were also bound by PDX1 in the adult islet in both mouse and human (bold gene in Figure 5F). These ductal specific genes are known to be involved in mitochondrial function (*ATP1B1* and *ATP1A1*), related to pancreatic cancer progression (*ACTN4, TPM1*, *CYR61, CLDN3*) (31, 32). In parallel, only 4 genes (*MEIS2, ISL1, PAX6,* and *SCGN*) were expressed in embryonic beta cells and bound by PDX1 in pancreas development in both mouse and human, which play important roles in pancreas development (33–37). *MEIS2, ISL1* and *PAX6* were also bound in the adult islet. Taken together, these results point towards a conserved developmental role of PDX1 primarily in the ductal and beta cell lineages.

## Discussion

We identified PDX1 genome-wide target genes *in vivo* in the developing pancreas of both mouse and human. The majority of PDX1 targets are conserved from developmental to adult stages and between mice and human, indicating PDX1 directs a core, developmentally and evolutionarily conserved gene program in the pancreatic islet. A subset of PDX1 targets were stage-specific and/or species-specific and could play important roles during pancreas development. By integrating ChIP-seq with single-cell RNA-seq data, we identified the temporal pattern of PDX1 targets expressed in each lineage. PDX1 surprisingly bound important acinar and ductal genes, from early development to adult islet in human pancreas, suggesting PDX1 as a keepsake of an embryonic pancreatic progenitor program in adult beta cells. We also identified potential PDX1 partners at different stages that likely synergize with PDX1 to execute its functions, which highlighted the dynamic changes in PDX1-coordinated networks in pancreas development.

### Identification of genome-wide PDX1 targets in mouse and human developing pancreas *in vivo*

The roles of PDX1 in the pancreas have been studied extensively due to its important roles in pancreas development and beta-cell function. Previous studies examined PDX1 genome-wide targets in human embryonic stem cell-derived pancreatic progenitors (16–18), which revealed the transcriptional networks regulated by PDX1 in pancreatic lineage development *in vitro*. By performing PDX1 ChIP-seq using human fetal pancreata at 14 weeks gestation and mouse embryonic pancreata at E13.5 and E15.5, we characterized the PDX1 cistrome of the developing pancreas. PDX1 directly regulated key transcription factors important in establishing pancreas lineages, including *PDX1*, *PTF1A, NKX6.1*, *HNF1B*, *ONECUT1*, and *GATA4*, which is consistent with previous *in vitro* studies. Moreover, we identified key signaling pathways important in pancreatic lineage establishment and expansion, including the WNT/beta-catenin signaling, epithelial-mesenchymal transition, Notch signaling, and Protein Kinase A pathways. The heterogeneity of the cells in the developing pancreas makes it difficult to identify PDX1 targets in individual cell types. By integrating PDX1 ChIP-seq with single-cell RNA-seq data, we revealed potential cell-lineage-specific PDX1 targets. It is surprising to identify the continuous binding of PDX1 on important acinar and ductal genes from early development to adult islet, revealing that the developmental role of PDX1 is conserved into the adult beta cell.

### PDX1 directs a core developmentally and evolutionarily conserved gene program in the pancreatic islet

During pancreas development, PDX1 is initially expressed in pancreatic progenitors to establish pancreatic lineages and then gradually enriched in pancreatic beta cells to maintain beta-cell function and identity (3, 4). The dramatic changes in PDX1 expression and function may be caused by changes in PDX1 binding. However, although part of PDX1 targets were stage-specific, as showed by Wang and colleagues (18), we found that PDX1 binding was largely stable from embryonic pancreas to adult islet in both mouse and human. The data indicated a core developmentally conserved PDX1 target program continues to exist in the pancreatic islet. Transcriptome analysis of pancreatic cells during development revealed that genes activated in the developing pancreas were largely conserved among vertebrates (38). Indeed, when comparing the PDX1 cistrome in mouse and human pancreas, we identified an evolutionarily conserved gene program regulated by PDX1. We found that there were 762 genes and 1509 genes bound by PDX1 in the developing pancreas and adult islets, respectively, of both mouse and human. The conserved genes play important roles in key functional pathways in pancreas development and function, including *Wnt/beta catenin signaling* at the developmental stage (39, 40), *Pi3K/AKT signaling* at adult stage (41), *protein kinase A signaling* at both developmental and adult stages (42).

### PDX1 interacts with other transcription factors and cofactors to regulate pancreas development

PDX1 have been found to act as both an activator and a repressor in some cell types. In hESC-pancreatic progenitors, PDX1 activates pancreatic genes and represses hepatic genes to establish pancreatic lineages (17). In adult beta cells, PDX1 activates key beta-cell functional genes and represses beta-cell disallowed genes to maintain normal beta function and identity (43). Our findings showed that PDX1 continuously bound the cis-regulatory regions of genes from developmental stages to adult stages, such as *Onecut1* and *Ptf1a*, which are expressed in PDX1+ pancreatic progenitors but not in PDX1+ beta cells or delta cells at adult stage. The data suggest that PDX1 activates *Onecut1* and *Ptf1a* in pancreatic progenitors but represses their expression in beta and delta cells. The activities of PDX1 depend in part on the cofactors it interacts with at different loci and/or in different cellular contexts. PDX1 may recruit different cofactors to activate or repress the expression of the same genes at different stages (44). Transcription factors interacting with PDX1 may also affect its downstream activities. We performed upstream regulator analysis and identified potential transcription factors and cofactors interacting with PDX1, which may coordinate with PDX1 to activate or repress its targets at different stages. The identified potential transcription factors and cofactors will provide valuable information for future studies to examine functionally-congruent or directly-interacting partners of PDX1 in the pancreas.

In summary, our study comprehensively examines PDX1 genome-wide targets in mouse and human developing pancreas and reveals a core, evolutionarily conserved gene program in the pancreatic islet. It enables future hypothesis-driven studies on the roles of PDX1 targets in mouse and human developing pancreas, provides the list of potential PDX1 partners that need to be present to allow for synergism of function, and sets a valuable foundation for evaluating the efficiency and improving the protocols for human stem cell differentiation to the beta-cell fate.

## Acknowledgements

JCR was supported by NIH F32-DK-089747. DES was supported by American Diabetes Association Junior Faculty Development Award (1-16-JDF086). DAS was supported by NIH grants R01DK105689 and R01 DK121175. We thank the University of Pennsylvania Diabetes Research Center (DRC) for the use of the Functional Genomics Core (P30-DK19525) and thank the Center for Applied Genomics at The Children’s Hospital of Philadelphia.

## Author contributions

JCR, DES designed and performed experiments, interpreted the results with input from DAS. XY wrote the manuscript, analyzed the data and interpreted the results. CL designed and performed experiments. JY performed experiments. RY, JK, GR performed the bioinformatic analysis, interpreted the data. KJW conceptualized the bioinformatic analysis and revised the manuscript. DAS conceptualized and supervised the project, wrote and edited the manuscript. DES conceptualized, analyzed the data, wrote/edited the manuscript.

---

**Supplemental Figure 1:**
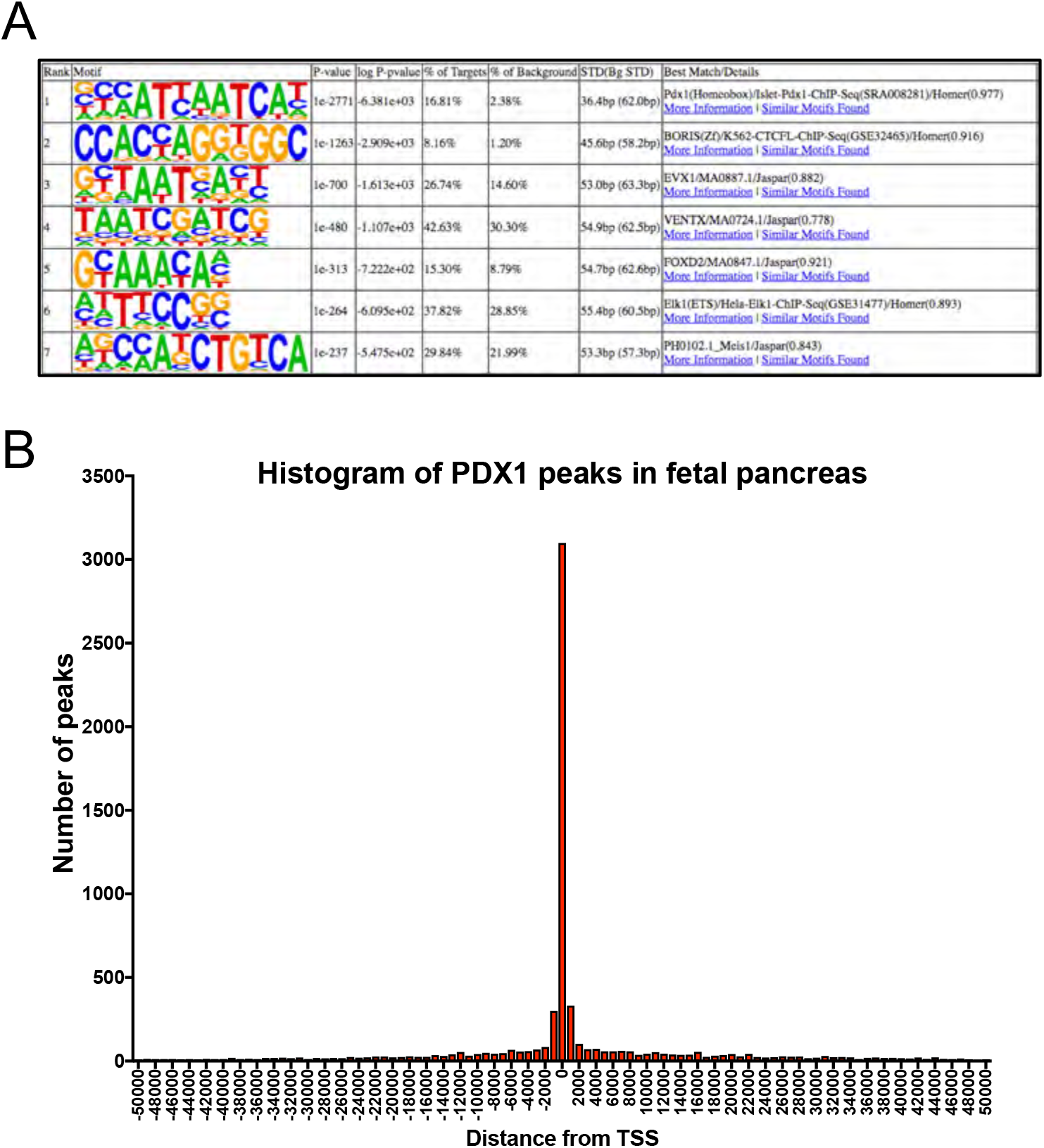
A. De novo motif analysis of PDX1 peaks in human fetal pancreas. B. Histogram distribution of PDX1 peaks in human fetal pancreas shows the majority of peaks are located within 2KB from the transcriptional starting site.

**Supplemental Figure 2:**
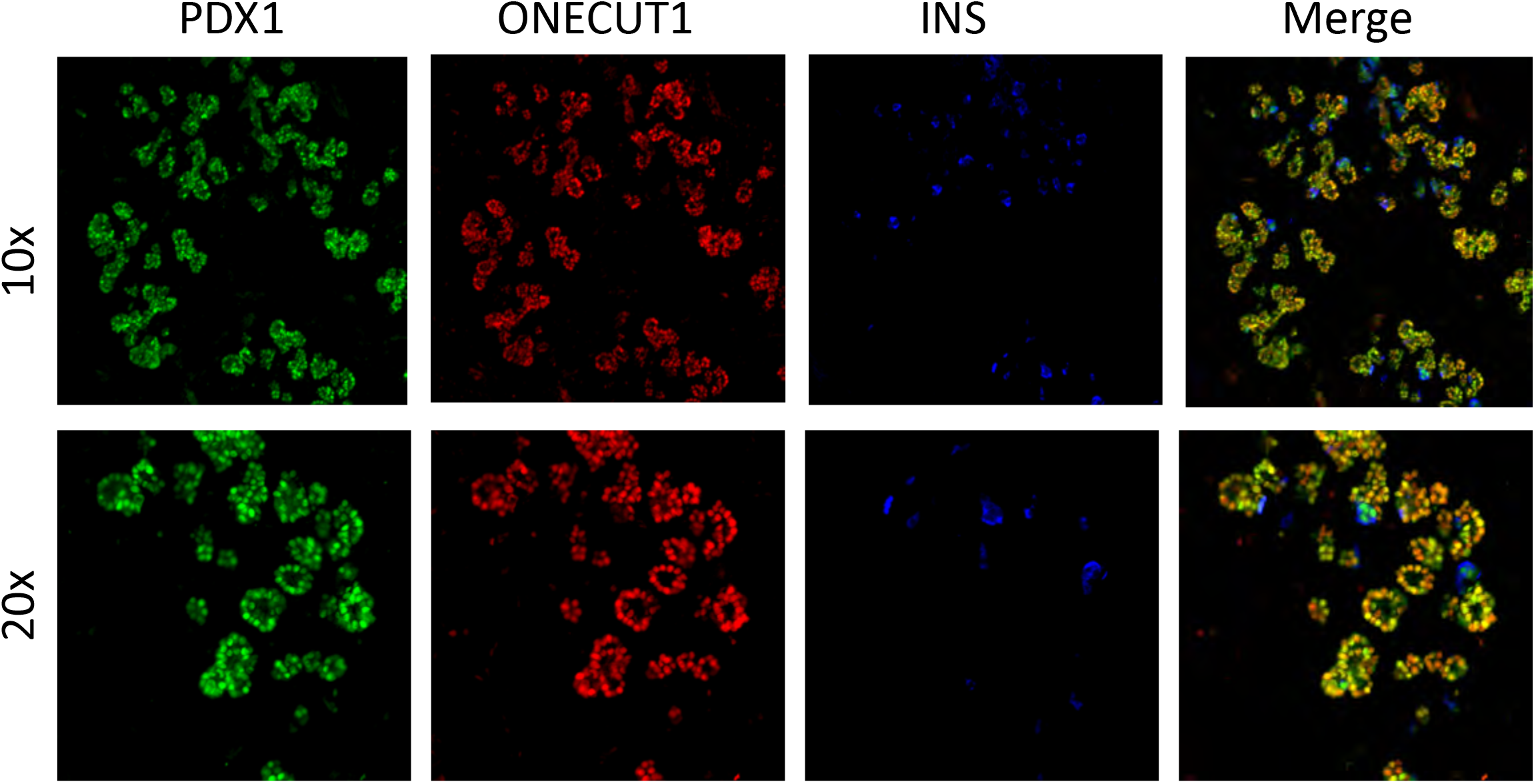
Immunofluorescence for PDX1, ONECUT1 and INS in human fetal pancreas at 14 weeks gestation shows that PDX1 is expressed in all pancreatic cells. INS+ cells express only PDX1, while the rest of the cells express both PDX1 and ONECUT1, showing these are acinar and ductal cells.

**Supplemental Figure 3:**
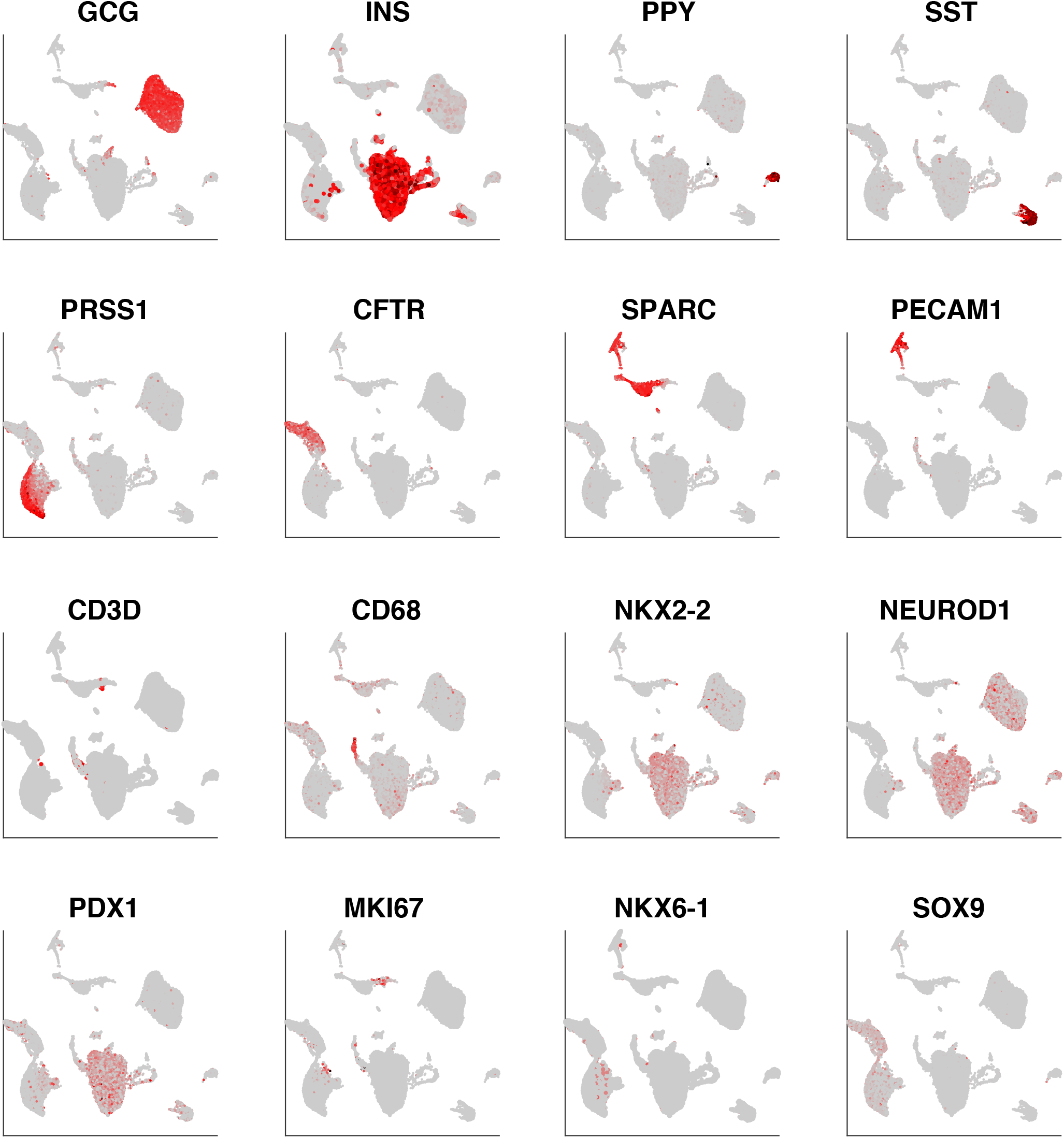
Representative expression of markers used to identify cell clusters in the UMAP plot of single-cell RNA-seq in human fetal pancreas.

**Supplemental Figure 4:**
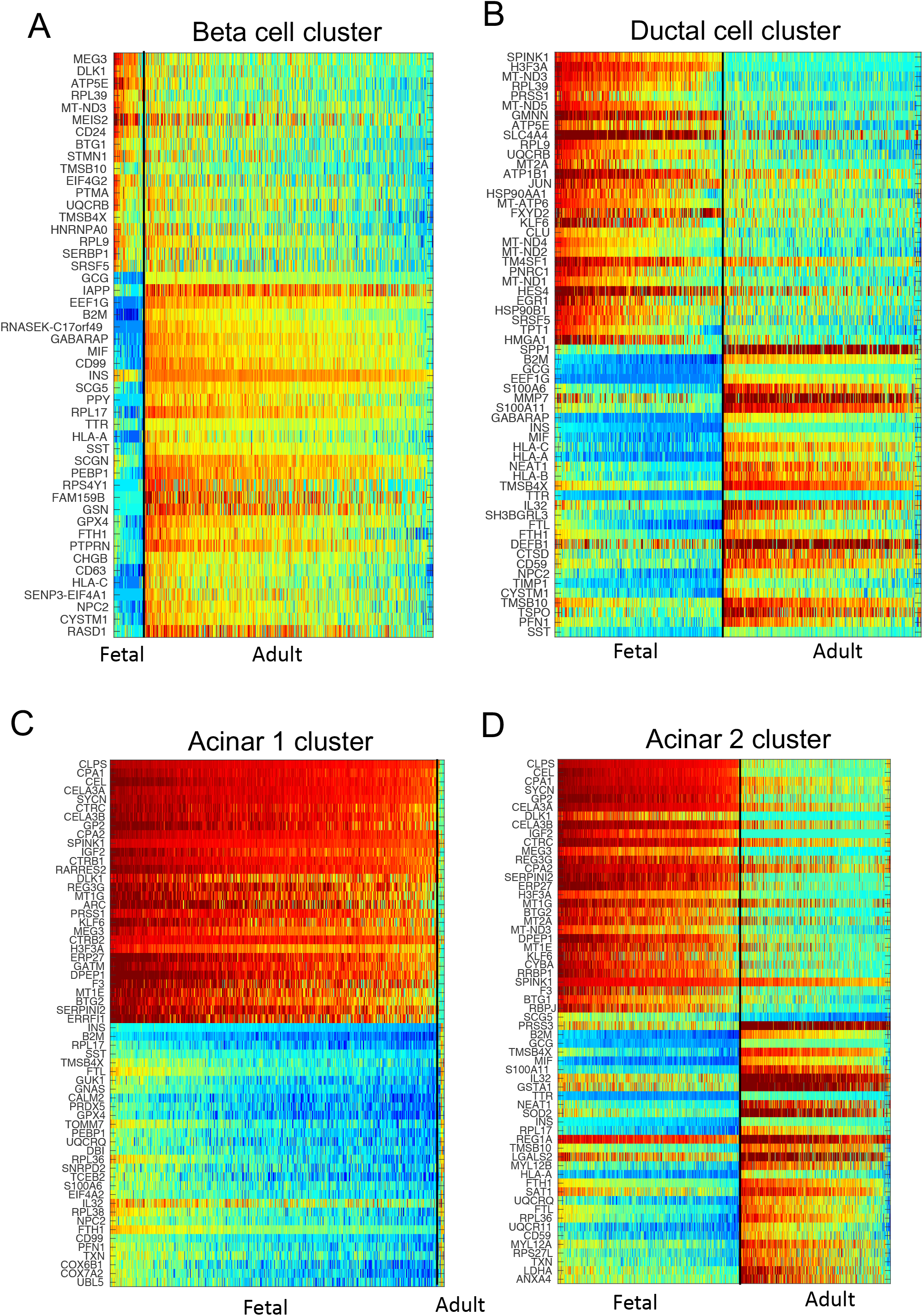
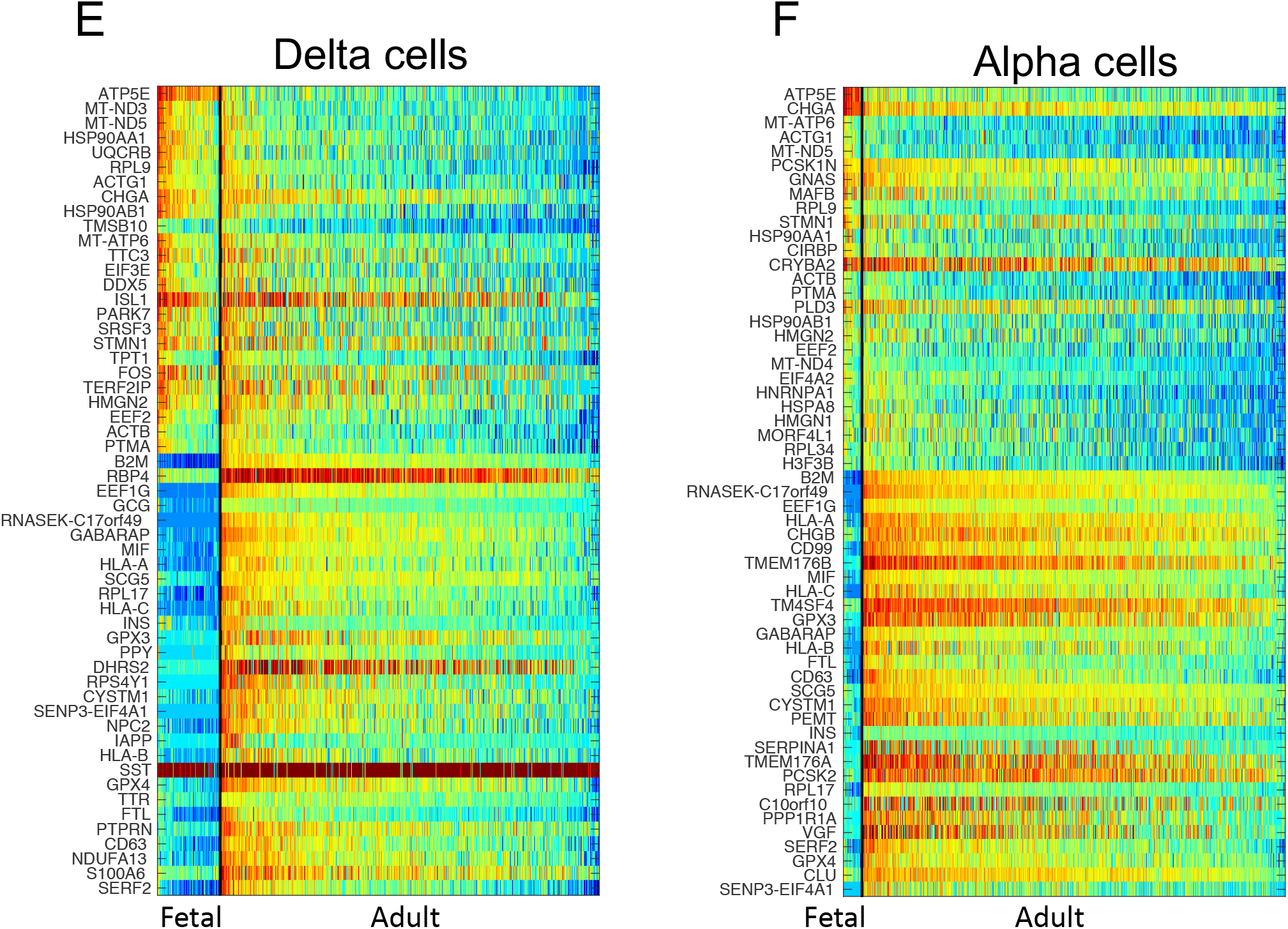
Heatmap showed differentially expressed genes between fetal and adult for each cell type. A. Genes differentially expressed between human fetal and adult beta cells. B. Genes differentially expressed between human fetal and adult ductal cells. C-D Genes differentially expressed between human fetal and adult acinar cells (Acinar 1 cluster mainly composed of fetal acinar cells, Acinar 2 cluster composed of both fetal and adult acinar cells). E. Genes differentially expressed between human fetal and adult delta cells. F. Genes differentially expressed between human fetal and adult alpha cells.

**Supplemental Figure 5:**
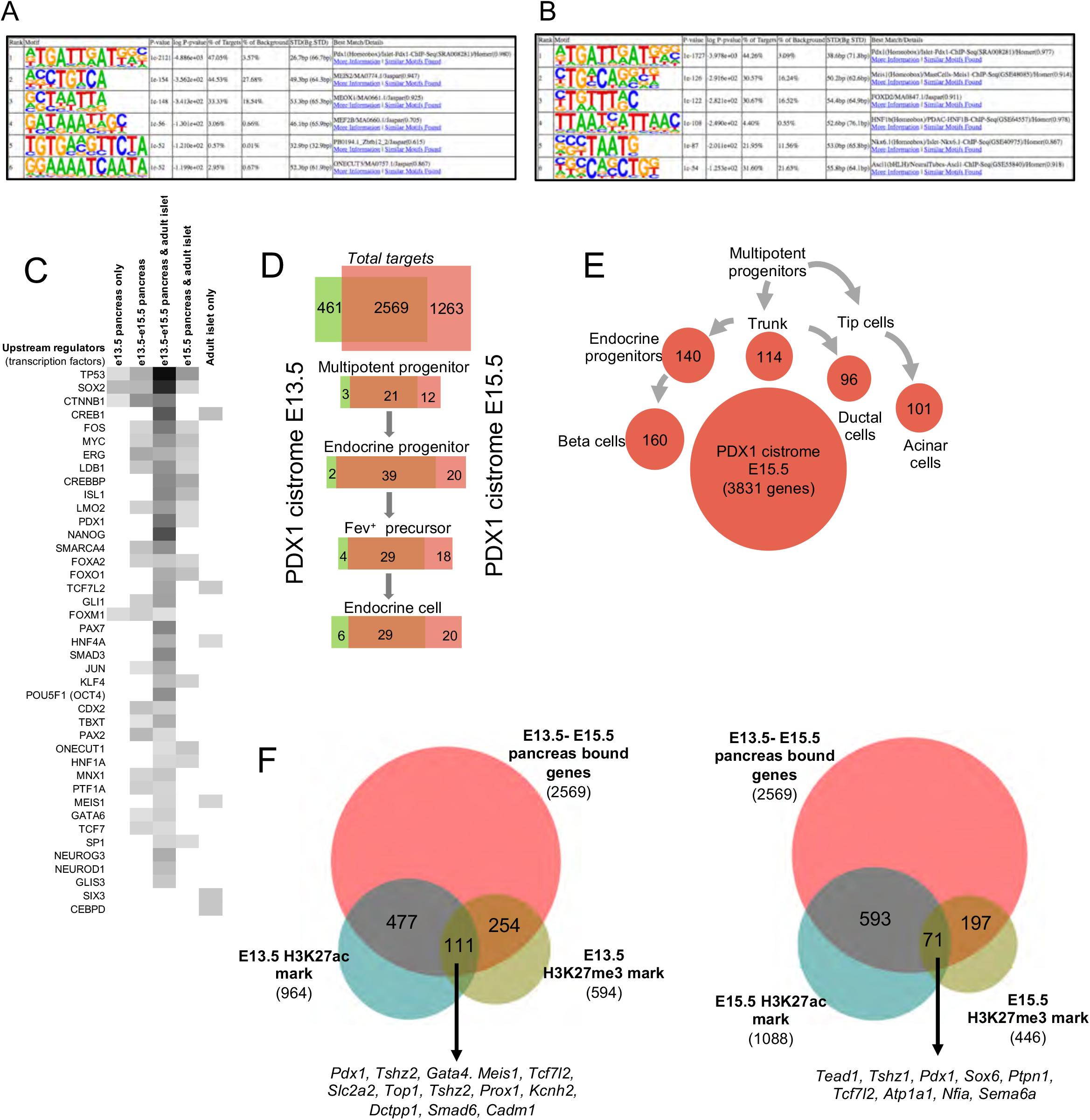
A-B. De novo motif analysis of PDX1 peaks in e13.5 (A) and e15.5(B) ChIP-seq data. C. Upstream regulator analysis of transcription factors predicted to directly regulate PDX1 bound genes in mouse developing pancreas and adult islet. D. Number of PDX1 bound genes expressed in different cell types in mouse developing pancreas at E13.5 and E15.5. E. PDX1 bound genes in E15.5 mouse pancreas and expressed in different cell types in mouse developing pancreas (based on comparison with Krentz, 2018). F. Histone modification analysis of PDX1 targets in mouse pancreas at E13.5 and E15.5.

**Supplemental Figure 6:**
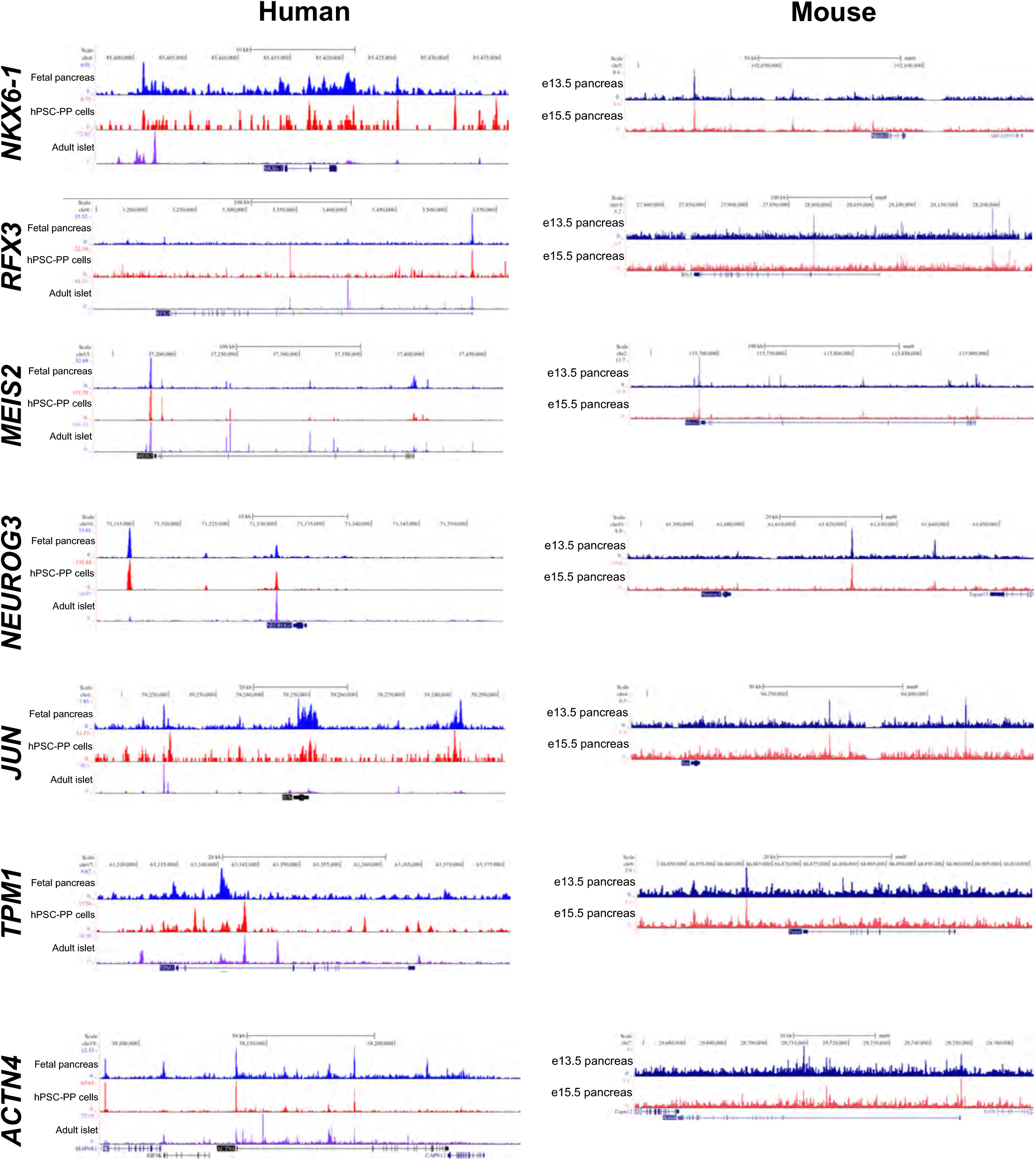
Pdx1 bound sites for selected genes (*NKX6.1, MEIS2, NEUROG3, RFX3, JUN, TPM1, ACTN4*)

## List of supplemental tables

**Table S1:** List of human fetal pancreas PDX1 peaks and bound genes

**Table S2:** List of hPSC-PP PDX1 peaks and bound genes

**Table S3:** List of human adult islet PDX1 peaks and bound genes

**Table S4:** Complete list of comparative canonical pathways gene ontology for genes bound by PDX1 in human pancreas data from Ingenuity IPA (by -log(p-value))

**Table S5:** Complete list of upstream regulators (transcription factors only) for genes bound by PDX1 in human pancreas data from Ingenuity IPA (by -log(p-value))

**Table S6:** List of differentially expressed genes among fetal pancreatic cells (FDR<0.01)

**Table S7:** List of differentially expressed gene among adult pancreatic cells (FDR<0.01) – analyzed data from Xin et al (Diabetes, 2018)

**Table S8:** List of genes bound by PDX1 in either fetal pancreas or in adult islet and expressed in either fetal pancreatic cells or adult pancreas cells

**Table S9:** List of E13.5 mouse pancreas PDX1 peaks and bound genes

**Table S10:** List of E15.5 mouse pancreas PDX1 peaks and bound genes

**Table S11:** List of mouse adult islet PDX1 peaks and bound genes

**Table S12:** Complete list of comparative canonical pathways gene ontology for genes bound by PDX1 in mouse pancreas data from Ingenuity IPA (by -log(p-value))

**Table S13:** Complete list of upstream regulators (transcription factors only) for genes bound by PDX1 in mouse pancreas data from Ingenuity IPA (by -log(p-value))

**Table S14:** List of genes bound by PDX1 in E13.5 or E15.5 pancreas in each type of pancreatic cells (based on data from Bastidas-Ponce et al, Development, 2019)

**Table S15:** List of genes bound by PDX1 in E15.5 pancreas in each type of pancreatic cells (based on data from Krentz et al, Stem Cell Reports, 2018)

**Table S16:** ChIP-seq for H3K27ac and H3K27me3 histone marks in E13.5 and E15.5 revealed active promoters and enhancers during mouse pancreas development

**Table S17:** List of common genes between human and mouse pancreas development and human and mouse adult islet

**Table S18:** Complete list of comparative canonical pathways gene ontology for genes bound by PDX1 in mouse and human pancreas data from Ingenuity IPA (by -log(p-value))

**Table S19:** Complete list of upstream regulators (transcription factors only) for genes bound by PDX1 in mouse and human pancreas data from Ingenuity IPA (by -log(p-value))

## References

1. IDF. The International Diabetes Federation; 2019.

2. Jennings RE, Berry AA, Strutt JP, Gerrard DT, and Hanley NA. Human pancreas development. Development. 2015;142(18):3126–37.

3. Pan FC, and Wright C. Pancreas organogenesis: from bud to plexus to gland. Dev Dyn. 2011;240(3):530–65.

4. Jennings RE, Scharfmann R, and Staels W. Transcription factors that shape the mammalian pancreas. Diabetologia. 2020;63(10):1974–80.

5. Guz Y, Montminy MR, Stein R, Leonard J, Gamer LW, Wright CV, et al. Expression of murine STF-1, a putative insulin gene transcription factor, in beta cells of pancreas, duodenal epithelium and pancreatic exocrine and endocrine progenitors during ontogeny. Development. 1995;121(1):11–8.

6. Jennings RE, Berry AA, Kirkwood-Wilson R, Roberts NA, Hearn T, Salisbury RJ, et al. Development of the human pancreas from foregut to endocrine commitment. Diabetes. 2013;62(10):3514–22.

7. Stanger BZ, Tanaka AJ, and Melton DA. Organ size is limited by the number of embryonic progenitor cells in the pancreas but not the liver. Nature. 2007;445(7130):886–91.

8. Roy N, Takeuchi KK, Ruggeri JM, Bailey P, Chang D, Li J, et al. PDX1 dynamically regulates pancreatic ductal adenocarcinoma initiation and maintenance. Genes Dev. 2016;30(24):2669–83.

9. Jonsson J, Carlsson L, Edlund T, and Edlund H. Insulin-promoter-factor 1 is required for pancreas development in mice. Nature. 1994;371(6498):606–9.

10. Offield MF, Jetton TL, Labosky PA, Ray M, Stein RW, Magnuson MA, et al. PDX-1 is required for pancreatic outgrowth and differentiation of the rostral duodenum. Development. 1996;122(3):983–95.

11. Stoffers DA, Zinkin NT, Stanojevic V, Clarke WL, and Habener JF. Pancreatic agenesis attributable to a single nucleotide deletion in the human IPF1 gene coding sequence. Nat Genet. 1997;15(1):106–10.

12. Stoffers DA, Ferrer J, Clarke WL, and Habener JF. Early-onset type-II diabetes mellitus (MODY4) linked to IPF1. Nat Genet. 1997;17(2):138–9.

13. Hani EH, Stoffers DA, Chèvre JC, Durand E, Stanojevic V, Dina C, et al. Defective mutations in the insulin promoter factor-1 (IPF-1) gene in late-onset type 2 diabetes mellitus. J Clin Invest. 1999;104(9):R41–8.

14. Oliver-Krasinski JM, Kasner MT, Yang J, Crutchlow MF, Rustgi AK, Kaestner KH, et al. The diabetes gene Pdx1 regulates the transcriptional network of pancreatic endocrine progenitor cells in mice. J Clin Invest. 2009;119(7):1888–98.

15. Henley KD, Stanescu DE, Kropp PA, Wright CVE, Won KJ, Stoffers DA, et al. Threshold-Dependent Cooperativity of Pdx1 and Oc1 in Pancreatic Progenitors Establishes Competency for Endocrine Differentiation and β-Cell Function. Cell Rep. 2016;15(12):2637–50.

16. Wang A, Yue F, Li Y, Xie R, Harper T, Patel NA, et al. Epigenetic priming of enhancers predicts developmental competence of hESC-derived endodermal lineage intermediates. Cell Stem Cell. 2015;16(4):386–99.

17. Teo AK, Tsuneyoshi N, Hoon S, Tan EK, Stanton LW, Wright CV, et al. PDX1 binds and represses hepatic genes to ensure robust pancreatic commitment in differentiating human embryonic stem cells. Stem Cell Reports. 2015;4(4):578–90.

18. Wang X, Sterr M, Burtscher I, Chen S, Hieronimus A, Machicao F, et al. Genome-wide analysis of PDX1 target genes in human pancreatic progenitors. Mol Metab. 2018;9:57–68.

19. Khoo C, Yang J, Weinrott SA, Kaestner KH, Naji A, Schug J, et al. Research resource: the pdx1 cistrome of pancreatic islets. Mol Endocrinol. 2012;26(3):521–33.

20. Hulsen T, de Vlieg J, and Alkema W. BioVenn - a web application for the comparison and visualization of biological lists using area-proportional Venn diagrams. BMC Genomics. 2008;9:488.

21. Xin Y, Dominguez Gutierrez G, Okamoto H, Kim J, Lee AH, Adler C, et al. Pseudotime Ordering of Single Human β-Cells Reveals States of Insulin Production and Unfolded Protein Response. Diabetes. 2018;67(9):1783–94.

22. Bastidas-Ponce A, Tritschler S, Dony L, Scheibner K, Tarquis-Medina M, Salinno C, et al. Comprehensive single cell mRNA profiling reveals a detailed roadmap for pancreatic endocrinogenesis. Development. 2019;146(12).

23. Krentz NAJ, Lee MYY, Xu EE, Sproul SLJ, Maslova A, Sasaki S, et al. Single-Cell Transcriptome Profiling of Mouse and hESC-Derived Pancreatic Progenitors. Stem Cell Reports. 2018;11(6):1551–64.

24. Morton JP, Timpson P, Karim SA, Ridgway RA, Athineos D, Doyle B, et al. Mutant p53 drives metastasis and overcomes growth arrest/senescence in pancreatic cancer. Proc Natl Acad Sci U S A. 2010;107(1):246–51.

25. Weissmueller S, Manchado E, Saborowski M, Morris JP, Wagenblast E, Davis CA, et al. Mutant p53 drives pancreatic cancer metastasis through cell-autonomous PDGF receptor β signaling. Cell. 2014;157(2):382–94.

26. Georgia S, Kanji M, and Bhushan A. DNMT1 represses p53 to maintain progenitor cell survival during pancreatic organogenesis. Genes Dev. 2013;27(4):372–7.

27. Schuijers J, Junker JP, Mokry M, Hatzis P, Koo BK, Sasselli V, et al. Ascl2 acts as an R-spondin/Wnt-responsive switch to control stemness in intestinal crypts. Cell Stem Cell. 2015;16(2):158–70.

28. Vanheer L, Schiavo A, Van Haele M, Haesen T, Janiszewski A, Chappell J, et al. bioRxiv (https://doi.org/10.1101/2020.09.23.310094); 2020.

29. Bastidas-Ponce A, Scheibner K, Lickert H, and Bakhti M. Cellular and molecular mechanisms coordinating pancreas development. Development. 2017;144(16):2873–88.

30. Benitez CM, Goodyer WR, and Kim SK. Deconstructing pancreas developmental biology. Cold Spring Harb Perspect Biol. 2012;4(6).

31. Abou-Kheir W, Mukherji D, Hadadeh O, Saleh E, Bahmad HF, Kanso M, et al. CYR61/CCN1 expression in resected pancreatic ductal adenocarcinoma: A retrospective pilot study of the interaction between the tumors and their surrounding microenvironment. Heliyon. 2020;6(5):e03842.

32. Westmoreland JJ, Drosos Y, Kelly J, Ye J, Means AL, Washington MK, et al. Dynamic distribution of claudin proteins in pancreatic epithelia undergoing morphogenesis or neoplastic transformation. Dev Dyn. 2012;241(3):583–94.

33. Ahlgren U, Pfaff SL, Jessell TM, Edlund T, and Edlund H. Independent requirement for ISL1 in formation of pancreatic mesenchyme and islet cells. Nature. 1997;385(6613):257–60.

34. Du A, Hunter CS, Murray J, Noble D, Cai CL, Evans SM, et al. Islet-1 is required for the maturation, proliferation, and survival of the endocrine pancreas. Diabetes. 2009;58(9):2059–69.

35. Malenczyk K, Szodorai E, Schnell R, Lubec G, Szabó G, Hökfelt T, et al. Secretagogin protects Pdx1 from proteasomal degradation to control a transcriptional program required for β cell specification. Mol Metab. 2018;14:108–20.

36. Ashery-Padan R, Zhou X, Marquardt T, Herrera P, Toube L, Berry A, et al. Conditional inactivation of Pax6 in the pancreas causes early onset of diabetes. Dev Biol. 2004;269(2):479–88.

37. Swift GH, Liu Y, Rose SD, Bischof LJ, Steelman S, Buchberg AM, et al. An endocrine-exocrine switch in the activity of the pancreatic homeodomain protein PDX1 through formation of a trimeric complex with PBX1b and MRG1 (MEIS2). Mol Cell Biol. 1998;18(9):5109–20.

38. Ramond C, Beydag-Tasöz BS, Azad A, van de Bunt M, Petersen MBK, Beer NL, et al. Understanding human fetal pancreas development using subpopulation sorting, RNA sequencing and single-cell profiling. Development. 2018;145(16).

39. Murtaugh LC. The what, where, when and how of Wnt/β-catenin signaling in pancreas development. Organogenesis. 2008;4(2):81–6.

40. Sharon N, Vanderhooft J, Straubhaar J, Mueller J, Chawla R, Zhou Q, et al. Wnt Signaling Separates the Progenitor and Endocrine Compartments during Pancreas Development. Cell Rep. 2019;27(8):2281–91.e5.

41. Kaneko K, Ueki K, Takahashi N, Hashimoto S, Okamoto M, Awazawa M, et al. Class IA phosphatidylinositol 3-kinase in pancreatic β cells controls insulin secretion by multiple mechanisms. Cell Metab. 2010;12(6):619–32.

42. Kaihara KA, Dickson LM, Jacobson DA, Tamarina N, Roe MW, Philipson LH, et al. β-Cell-specific protein kinase A activation enhances the efficiency of glucose control by increasing acute-phase insulin secretion. Diabetes. 2013;62(5):1527–36.

43. Gao T, McKenna B, Li C, Reichert M, Nguyen J, Singh T, et al. Pdx1 maintains β cell identity and function by repressing an α cell program. Cell Metab. 2014;19(2):259–71.

44. Spaeth JM, Walker EM, and Stein R. Impact of Pdx1-associated chromatin modifiers on islet β-cells. Diabetes Obes Metab. 2016;18 Suppl 1:123–7.

